# A high-throughput assay for directly monitoring nucleolar rRNA biogenesis

**DOI:** 10.1101/2021.07.19.452935

**Authors:** Carson J. Bryant, Mason A. McCool, Laura Abriola, Yulia V. Surovtseva, Susan J. Baserga

## Abstract

Studies of the regulation of nucleolar function are critical for ascertaining clearer insights into the basic biological underpinnings of ribosome biogenesis, and for future development of therapeutics to treat cancer and ribosomopathies. A number of high-throughput primary assays based on morphological alterations of the nucleolus can indirectly identify hits affecting ribosome biogenesis. However, there is a need for a more direct high-throughput assay for nucleolar function to further evaluate hits. Previous reports have monitored nucleolar RNA biogenesis using 5-ethynyl uridine (5-EU) in low-throughput. We report a miniaturized, high-throughput 5-EU assay for nucleolar function which enables specific calculation of nucleolar rRNA biogenesis inhibition, based on co-staining of the nucleolar protein fibrillarin (FBL). The assay utilizes two siRNA controls, a negative non-targeting siRNA control (siNT) and a positive siRNA control targeting *POLR1A* (siPOLR1A), and specifically quantifies median 5-EU signal within nucleoli. Maximum nuclear 5-EU signal can also be used to monitor the effects of putative small molecule inhibitors of RNAP1, like BMH-21, or other treatment conditions that cause FBL dissociation. We validate the 5-EU assay on 68 predominately nucleolar hits from a high-throughput primary screen, showing that 58/68 hits significantly inhibit nucleolar rRNA biogenesis. Our new method establishes direct quantification of nucleolar function in high-throughput, facilitating closer study of ribosome biogenesis in health and disease.

## Introduction

Cells of all organisms manufacture mature ribosomes, the core machinery of protein translation, through a process known as ribosome biogenesis (RB) (reviewed in Woolford and Baserga, 2013; Aubert et al., 2018). In eukaryotic cells, the first steps of RB occur in the nucleolus, a membraneless nuclear organelle discovered in the 1830’s (reviewed in Pederson, 2011; Correll et al., 2019; Lafontaine et al., 2020), where RNA Polymerase 1 (RNAP1) transcribes the primary pre-ribosomal RNA (pre-rRNA) precursor (reviewed in Moss et al., 2007; Sharifi and Bierhoff, 2018; Panov et al., 2021). Subsequently, a series of RNA processing and modification steps transpire, largely within the nucleolus, to create the mature cytoplasmic 18S, 5.8S, and 28S rRNA molecules in human cells (Henras et al., 2015; Sloan et al., 2017). Ribosomal proteins (RPs) bind (pre-)rRNA substrates in a hierarchical progression throughout this maturation process, bolstering the stability of the nascent transcript by chaperoning its folding away from incorrect, energetically-minimized conformations (O’Donohue et al., 2010; de la Cruz et al., 2015). Dysregulation of RB, and particularly of RNAP1 transcription, is a causative factor in a myriad of human disease states, including cancer (Derenzini et al., 2017; Bustelo and Dosil, 2018; Pelletier et al., 2018; Low et al., 2019; Penzo et al., 2019; Harold et al., 2021), ageing (Hein et al., 2012; Sharifi and Bierhoff, 2018), and rare diseases called ribosomopathies (Yelick and Trainor, 2015; Aspesi and Ellis, 2019; Farley-Barnes et al., 2019).

Given the importance of nucleolar function in human health and disease, the creation of more robust tools for measuring rRNA biogenesis within the nucleolus is essential for understanding the basic biological mechanisms through which RB can be regulated, as well as for developing next-generation small molecule or biologic therapeutics. In the past decade, a cadre of studies using high-throughput screening (HTS) have elucidated novel mechanisms through which human RB is regulated (Tafforeau et al., 2013; Badertscher et al., 2015; Farley-Barnes et al., 2018; Ogawa et al., 2021); several candidate therapeutics targeting the nucleolus have also been discovered with HTS chemical library or natural product campaigns (Drygin et al., 2011; Peltonen et al., 2014b; Scull et al., 2019; Ferreira et al., 2020; Kirsch et al., 2020). While several HTS modalities for monitoring nucleolar form and morphology have been described (Farley-Barnes et al., 2018; He et al., 2018; Stamatopoulou et al., 2018), none of these platforms directly measure the nucleolus’ rRNA biosynthetic function. To date, the lack of a direct high-throughput assay for nucleolar rRNA biogenesis constrains researchers’ ability to select for and validate the most promising candidate regulators of RB.

To monitor nucleolar function in a high-throughput manner, we sought to adapt a 5-ethynyl uridine (5-EU) assay for nucleolar rRNA biogenesis to an accessible, miniaturized format. The 5-EU assay has been successfully used to quantify changes in nucleolar transcriptional activity by several other groups in a variety of systems including human tissue culture cells (Bai et al., 2013; Lafita-Navarro et al., 2016; Cheng et al., 2017; Hayashi et al., 2017; Calo et al., 2018; Hayashi et al., 2018; Rossetti et al., 2018; Stamatopoulou et al., 2018; Dong et al., 2021), primary neurons (Slomnicki et al., 2018), porcine fetal fibroblasts (Lin et al., 2020), *Drosophila melanogaster* ovarian stem cells (Mikhaleva et al., 2019), and plant seedlings (Dvořáčková and Fajkus, 2018; Hayashi and Matsunaga, 2019). A key limitation in almost all of these studies is that total cellular or total nuclear 5-EU is quantified, rather than solely nucleolar 5-EU. Because only nucleolar signal corresponds to biogenesis of the primary pre-rRNA, quantifying total 5-EU leads to increased background from nascent transcription by RNA Polymerases besides RNAP1. Additionally, the computational methods used for image segmentation and quantification have varied widely and include custom MATLAB scripts, manual definition of regions-of-interest in ImageJ, and image multiplication in Adobe Photoshop, further limiting assay accessibility and reproducibility across research groups.

To improve upon these limitations, we present a miniaturized, high-throughput-ready 5-EU assay that selectively measures nucleolar rRNA biogenesis by co-staining for the nucleolar protein fibrillarin (FBL). In addition, we provide an analysis pipeline for the open-source image analysis software CellProfiler (McQuin et al., 2018) that provides a facile and reproducible framework for quantifying nucleolar 5-EU levels. We validate our assay by depleting 68 known RB factors including core RNAP1 machinery, assembly factors, and RPs, demonstrating robust and reproducible results for specifically measuring nucleolar rRNA biogenesis. Overall, our miniaturized 5-EU assay expands the dimensionality of HTS experiments studying the nucleolus, and will accelerate the discovery of novel RB regulators and targeted therapeutics.

## Results

### A high-content assay to quantify nucleolar rRNA biogenesis

In order to achieve specific quantification of nucleolar rRNA biogenesis, we introduced a 5-EU labeling step into our previously established screening platform for counting nucleolar number (Farley-Barnes et al., 2018), which utilizes CellProfiler (McQuin et al., 2018) to segment nuclei and nucleoli in images of cells immunofluorescently stained for DNA and the nucleolar protein fibrillarin (FBL) (**Figure 1A**). In our new protocol, MCF10A breast epithelial cells are reverse-transfected with siRNA duplexes for 72 h. For one hour following the transfection period, the cells are treated with 1 mM 5-EU, which is incorporated into nascent transcripts. Since the bulk of cellular transcription occurs in the nucleolus, most of the 5-EU label is incorporated into nucleolar nascent pre-rRNA (**Figure 1A**). The cells are fixed and immunofluorescently stained for DNA and FBL, after which nascent RNA is visualized *in situ* by performing a bio-orthogonal click reaction to covalently label the 5-EU alkyne moiety with an azide fluorophore (AF488 azide) (**Figure 1A**). The cells are then imaged and analyzed with CellProfiler to specifically quantify nucleolar rRNA biogenesis across all control and unknown wells. CellProfiler is known for its ease-of-use and modular adaptability (Stoter et al., 2019; Dobson et al., 2021), making it suitable for inclusion in a standardized, broadly accessible protocol.

**Figure 1.**
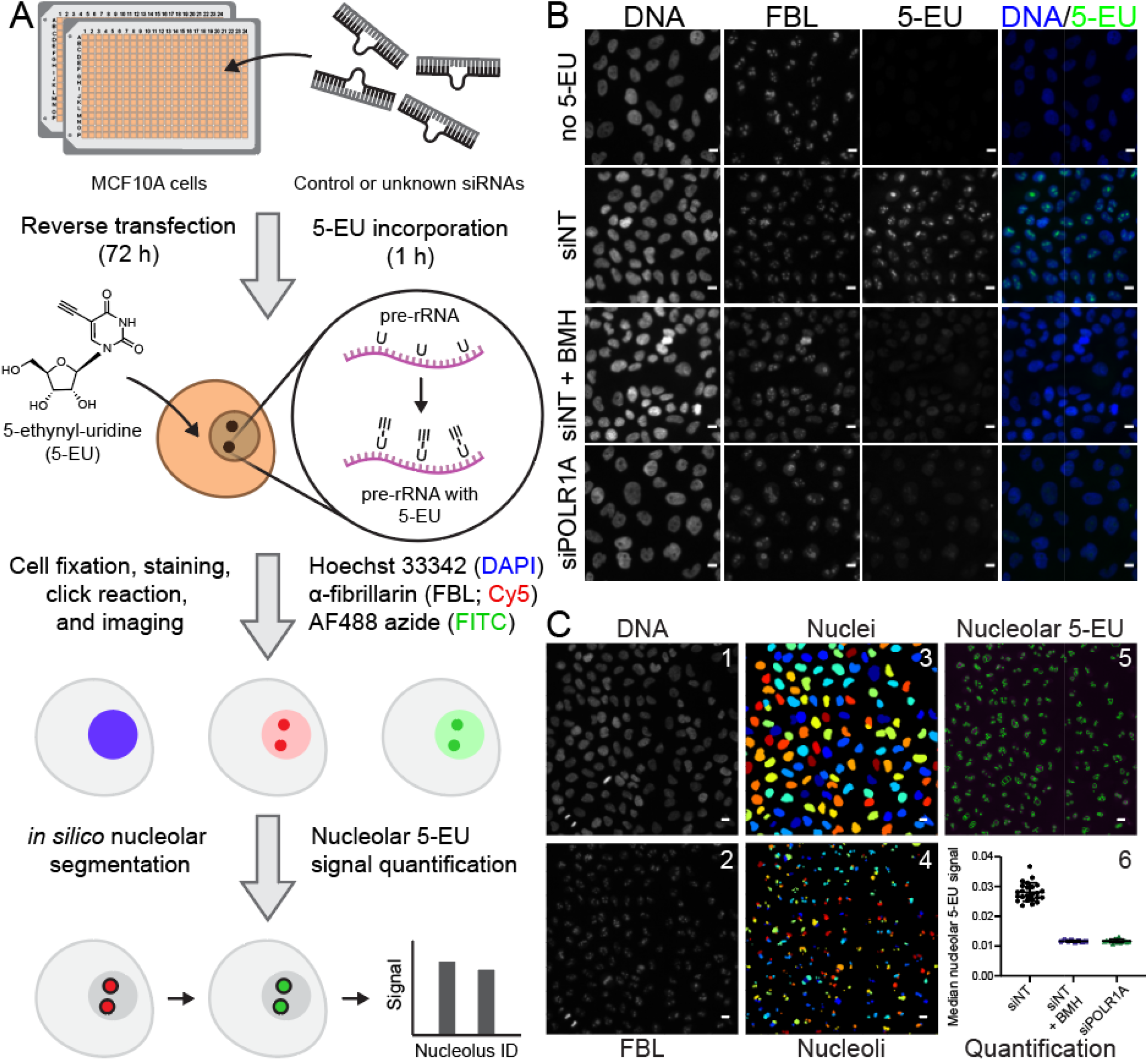
A high-throughput assay for nucleolar rRNA biogenesis using 5-ethynyl uridine (5-EU). **A.** Schematic of the 5-EU assay protocol. MCF10A cells are reverse-transfected in 384-well imaging plates with control or unknown siRNAs for 72 h. Following target depletion, 5-EU is incorporated into nascent RNA transcripts for 1 h, with the majority of label incorporated into nascent pre-ribosomal RNA (pre-rRNA). Treated cells are fixed and stained for DNA (Hoechst 33342, DAPI channel) and the nucleolar protein fibrillarin (FBL, Cy5 channel). 5-EU in nascent transcripts is conjugated to an azide fluorophore (AF488 azide, FITC channel) via a copper-catalyzed click reaction. After fluorescent imaging, cell nuclei and nucleoli are segmented *in silico* with CellProfiler, and nucleolar-specific 5-EU signal is quantified for each nucleolus object identified. **B.** RNAP1 inhibition specifically inhibits nucleolar 5-EU incorporation. No 5-EU, experiment without 1 h 5-EU incorporation. Treatment with a non-targeting siRNA (siNT) leads to high 5-EU signal within the nucleolus and moderate nucleoplasmic background signal. Acute treatment with BMH-21 (siNT + BMH) or siRNA-mediated depletion of POLR1A (siPOLR1A) decreases nucleolar 5-EU signal, although nucleoplasmic background remains. DNA (Hoechst staining), FBL (fibrillarin staining), 5-EU (5-EU staining), DNA/5-EU (combined Hoechst and EU staining). Scale bars, 10 μm. **C.** Schematic of CellProfiler segmentation and nucleolar 5-EU quantification. Panels 1 and 2, raw images of DNA and FBL staining. Panels 3 and 4, nuclei or nucleoli segmented by CellProfiler from DNA or FBL staining, respectively. Rainbow coloring identifies object number. Panel 5, overlay of segmented nucleoli (green) on top of 5-EU staining (magenta). Panel 6, quantification of median nucleolar 5-EU signal for nucleoli in cells treated with siNT, siNT and BMH-21, or siPOLR1A. n = 24, 8, or 16 wells, respectively. Scale bars, 10 μm.

We optimized our 5-EU assay to use a non-targeting siRNA as a negative control (siNT), and an siRNA targeting *POLR1A*, the largest subunit of RNAP1 also known as RPA194, as a positive control (siPOLR1A) (**Figure 1B**). RNAP1 inhibition by POLR1A depletion strongly reduces the nucleolar 5-EU signal to a degree consistent with acute treatment with BMH-21, a potent small molecule inhibitor of RNAP1 (Peltonen et al., 2014a; Peltonen et al., 2014b) (**Figure 1B-C**, compare siNT to siNT + BMH and siPOLR1A). However, it is clear that residual nucleoplasmic 5-EU signal remains even after RNAP1 inhibition (**Figure 1B**, siNT + BMH and siPOLR1A), emphasizing the importance of only quantifying 5-EU staining within the nucleolus via FBL co-staining.

To achieve nucleolar 5-EU quantification during analysis, images of DNA and FBL staining (**Figure 1C**, panels 1 and 2) were first used to segment nuclei and nucleoli by CellProfiler (**Figure 1C**, panels 3 and 4), respectively. Then, the median 5-EU signal within each nucleolus was measured (**Figure 1C**, panel 5), enabling aggregate quantification analysis per treatment condition across every nucleolus within each well (**Figure 1C**, panel 6). Final calculation of mean signals, percent inhibitions (by normalization to the negative and positive controls), and screening statistics including signal-to-background (S/B) and Z’ factor can be carried out in any standard data analysis software that can import the CellProfiler output CSV files, such as Microsoft Excel, JMP, R, or Python pandas.

### Optimization of the 5-EU assay for a miniaturized 384-well plate format

To adapt the 5-EU assay for use in high-throughput, we developed and optimized an optional 5-EU module that integrates into our existing nucleolar number screening platform. We first investigated the assay in MCF10A cells in the absence of siRNA knockdown or FBL co-staining, using the potent RNAP1 inhibitor BMH-21 or DMSO vehicle as positive or negative controls, respectively. For the first optimization experiments without FBL co-staining, median or maximum nuclear 5-EU signal was measured. We hypothesized that maximum nuclear 5-EU signal should track nucleolar function more accurately than the median, since a larger difference in the maximum value should be observed after RNAP1 inhibition. However, both metrics should decrease significantly upon BMH-21 treatment. Based on the original 5-EU method publication (Jao and Salic, 2008), we chose to label cells with 5-EU for 1 h, striking a balance between signal levels and incorporation time. By varying the 5-EU treatment concentration and click reaction time in wells treated without or with BMH-21, we discovered that treatment with 1 mM 5-EU for 1 h, followed by a 30 min click reaction was optimal (**Figure 2A**). Specifically, these conditions achieved the highest S/B ratio for the controls for each metric (**Figure 2B**).

**Figure 2.**
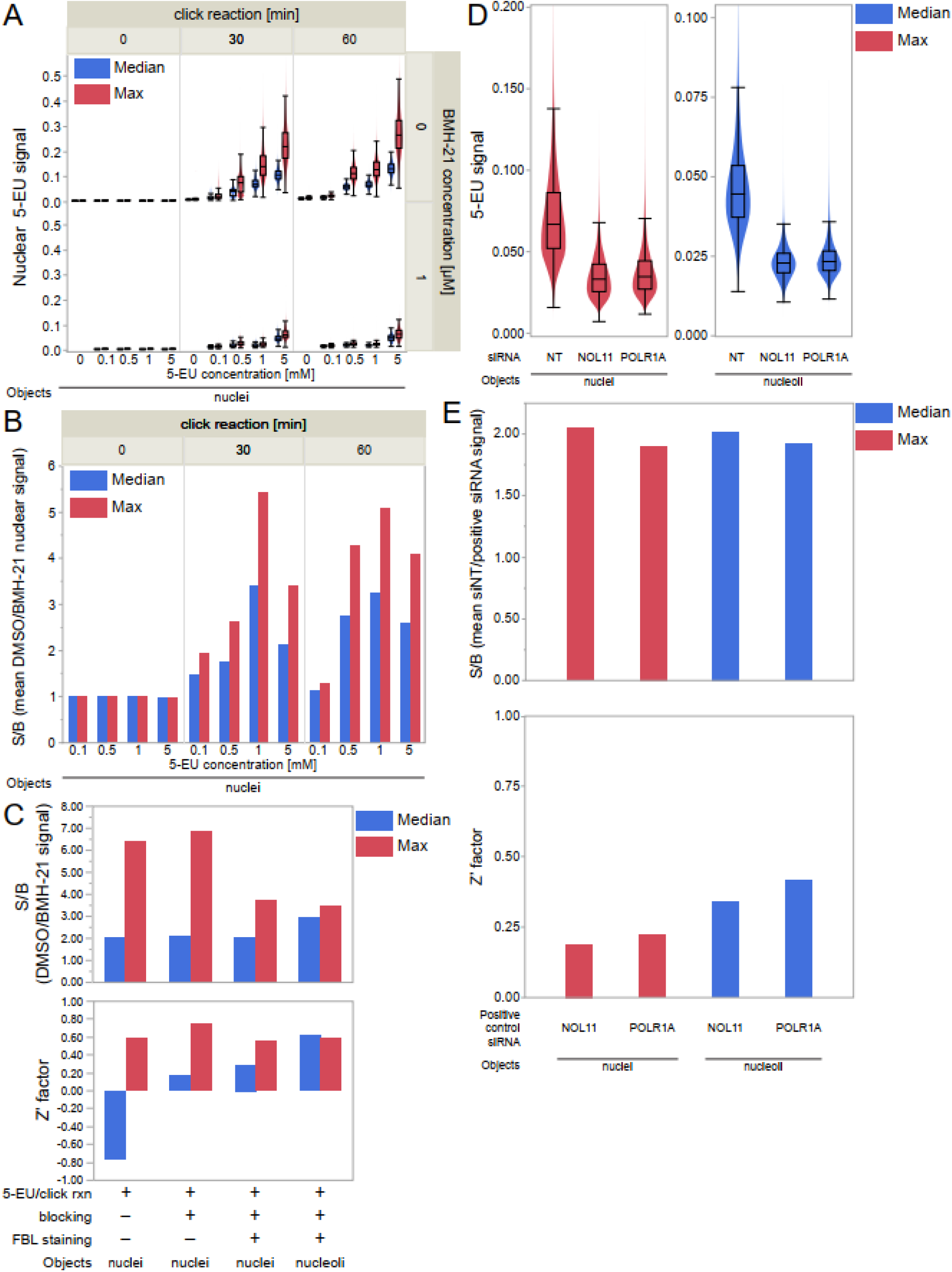
Optimization of the miniaturized 5-EU assay for nucleolar rRNA biogenesis. **A**. Median (blue) or maximum (red) nuclear 5-EU signal for cells treated across a range of 5-EU concentrations, click reaction times, and BMH-21 concentrations. n ≥ 20,000 cells per condition. **B.** Control signal-to-background (S/B) ratios for treatment conditions in Panel A. Control S/B is calculated as the ratio of mean DMSO-treated nuclear 5-EU signal divided by mean BMH-21-treated nuclear 5-EU signal, for each combination of 5-EU concentration and click reaction time. Median nuclear 5-EU signal (blue), maximum nuclear 5-EU signal (red). **C.** Control S/B and Z’ factor values for nuclei or nucleoli objects with only 5-EU visualization, 5-EU plus blocking with 10% (v/v) FBS/PBS, or 5-EU plus blocking and FBL co-staining. Median 5-EU signal (blue), maximum 5-EU signal (red). **D.** Maximum nuclear 5-EU signal (red) or median nucleolar 5-EU signal (blue) for cells treated with siNT, siNOL11, or siPOLR1A. n ≥ 130,000 cells per siRNA. **E.** Control S/B and Z’ factor values for cells (from Panel D) treated with siNOL11 or siPOLR1A as the positive control. Control S/B is calculated as the ratio of mean siNT-treated 5-EU signal divided by mean siNOL11- or siPOLR1A-treated 5-EU signal. Maximum nuclear 5-EU signal (red), median nucleolar 5-EU signal (blue).

Next, we introduced steps to enable nucleolar segmentation including blocking with a 10% (v/v) FBS/PBS solution and immunofluorescent staining for FBL. We utilized blocking and staining parameters that were previously optimized for our original screening platform (Farley-Barnes et al., 2018). Using BMH-21, we found the highest control S/B and Z’ factor occurred when measuring maximum 5-EU signal in nuclei that had been blocked but not stained for FBL (**Figure 2C**, second group). To specifically quantify nucleolar 5-EU incorporation, we also measured median nucleolar 5-EU signal. We selected the median metric because, compared to the maximum, it is more robust to outliers that may occur from staining artifacts or other abnormalities. When segmenting nucleoli, a comparable Z’ factor was achieved when measuring median 5-EU signal in nucleoli (**Figure 2C**, fourth group).

Interestingly, we also noted that acute treatment with BMH-21 during our 1 h 15 min treatment period caused increased nucleoplasmic FBL staining, presumably from FBL dissociation following RNAP1 inhibition (**Figure 1B**, siNT + BMH, FBL panel). We caution that the accuracy of nucleolar segmentation should be closely monitored if BMH-21 or another potent RNAP1 inhibitor that causes FBL dissociation is used. Aberrancies in FBL staining could lead to inaccurate segmentation, affecting results obtained by calculating median nucleolar signal. In these situations, maximum nuclear signal can be monitored.

In the final phase of optimization, we studied how siRNA knockdown of known RB factors affected nuclear and nucleolar 5-EU signal. We chose to deplete NOL11, a small subunit processome factor critical for pre-rRNA transcription (Freed et al., 2012), or POLR1A, the largest subunit of the RNAP1 complex, as positive controls. Compared to treatment with siNT, depletion of NOL11 or POLR1A decreased maximum nuclear signal and median nucleolar signal by roughly 50% in each case (**Figure 2D**), corresponding to control S/B values of 1.9-2.0 for each control (**Figure 2E**, top). However, measuring median nucleolar signal had lower object-to-object variability, resulting in more favorable Z’ factors than when measuring maximum nuclear signal (**Figure 2E**, bottom). Thus, both NOL11 and POLR1A are excellent positive controls for inhibiting nucleolar rRNA biogenesis in the 5-EU assay, when median nucleolar signal is measured. In follow-up validation studies (see below), we confirmed that measuring the median nucleolar 5-EU signal provides the most robust Z’ factors, despite the nucleolar 5-EU standard deviation metric having a higher control S/B ratio (**Supplementary Figure 1**). From these results, we recommend measuring maximum nuclear 5-EU for cell line optimization studies using very potent small molecule inhibitors of nucleolar function, such as BMH-21. We also conclude that measuring median nucleolar 5-EU signal, which corresponds only to nucleolar rRNA biogenesis, is the optimal 5-EU assay endpoint.

### Validation of the high-throughput 5-EU assay on 68 known ribosome biogenesis factors

After optimization, we validated the high-throughput 5-EU assay using a subset of 68 previously studied RB factors, including RPs and assembly factors for both ribosomal subunits, as well as core RNAP1 machinery and drivers of transcription such as MYC (**Figure 3A, Supplementary Table 1**). We depleted each RB factor over 72 h using siRNA pools in accordance with our protocol, performing the assay in biological triplicate to ensure reproducibility. Strikingly, we found that depletion of 58/68 biogenesis factors led to a significant (≥ 50%) inhibition of nucleolar 5-EU signal after standardization to the controls (**Figure 3A**). Images of the assay controls illustrate typical signal levels observed for the negative control siNT, set at 0% inhibition, and the positive control siPOLR1A, set at 100% inhibition (**Figure 3B**, siNT and siPOLR1A). Furthermore, images from the RB factors tested demonstrate the sensitivity of the assay to RNAP1 inhibition, from extreme effects above 100% inhibition (*e.g.* siMYC) to more moderate inhibitory effects (*e.g.* siTRMT112) (**Figure 3B**). Full results from the assay validation are presented in **Figure 3C** and **Supplementary Table 1**.

**Figure 3.**
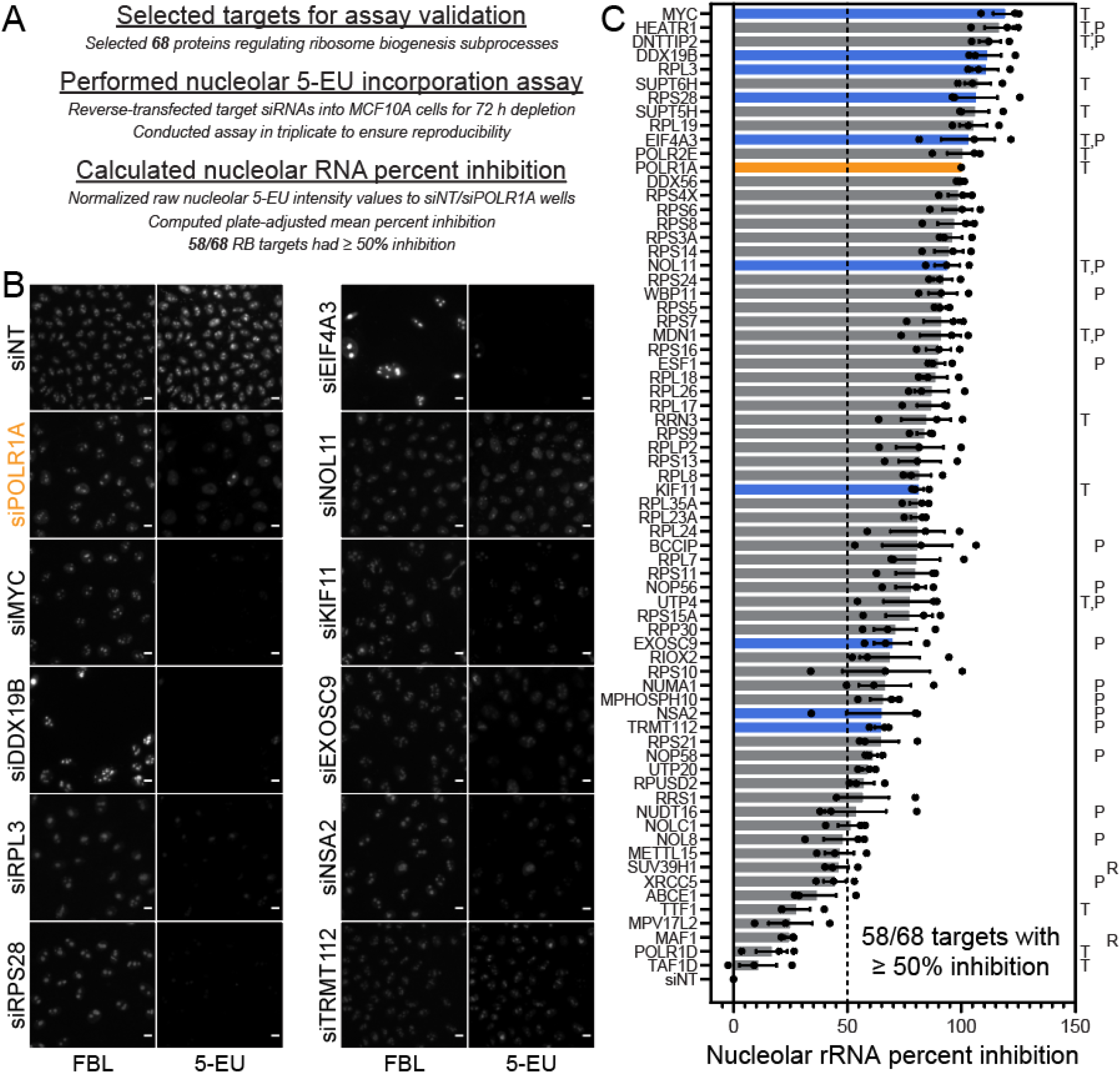
Validation of the 5-EU assay for nucleolar rRNA biogenesis on 68 known ribosome biogenesis factors. **A.** Outline of assay validation experiments. 68 proteins known to regulate ribosome biogenesis subprocesses, including RNAP1 transcription and pre-rRNA processing, modification, or stability, were selected for assay validation. The 5-EU assay was performed on cells depleted of these factors in biological triplicate, as described. **B.** Representative images of FBL staining and 5-EU visualization for cells treated with siNT (negative control), siPOLR1A (positive control, orange), or a subset of siRNAs targeting known RB factors. **C.** Nucleolar rRNA percent inhibition values for cells depleted of each known RB factor. Black dots, individual percent inhibition values for one biological replicate. Solid bars, mean percent inhibition (n = 3). Orange bar, PO R1A positive control (percent inhibition = 100%). Blue bars, RB factors illustrated in Panel B. Letters to right indicate factors involved in RNAP1 transcription (T), pre-rRNA processing (P), or transcription repression (R).

## Discussion

More precise, accessible methods for the study of nucleolar function are critical for illuminating novel ribosome biogenesis regulators and next-generation therapeutics for human disease states including cancer, aging, and rare ribosomopathies. Here, we developed an HT-ready, image-based assay that selectively measures nucleolar rRNA biogenesis in MCF10A breast epithelial cells. Building upon previous HTS techniques, we combined FBL staining of nucleoli and 5-EU incorporation into nascent RNA to measure only the 5-EU signal corresponding to nucleoli. We optimized the parameters of this assay using both small molecule inhibition (BMH-21) and acute siRNA depletion of essential RNAP1 transcription machinery (POLR1A and NOL11). Our detailed assay framework can be applied to studies of novel RNAP1 drug inhibitors and cellular regulators of nucleolar rRNA biogenesis, with the potential for adaptation to a variety of cell types. Our assay will increase the dimensionality and efficiency of future HTS campaigns focused on the nucleolus, accelerating the discovery of novel modulators of nucleolar function,

After optimizing the 5-EU assay for a miniaturized format, we validated its utility on 68 known RB factors including core RNAP1 components, small (pre-40S) or large (pre-60S) ribosomal subunit-specific processing and assembly factors, pre-rRNA modification factors, and RPs. As expected, all RB factors had a percent inhibition value greater than 0%. While a wide range of percent inhibition values were observed, 58/68 factors (85.5%) had a mean percent inhibition of at least 50%, signaling that the 5-EU assay robustly reports depletion conditions that interrupt nucleolar rRNA biogenesis.

Strikingly, we observed 11 targets that resulted in stronger nucleolar 5-EU inhibition than the positive control, POLR1A; consistent with a mean percent inhibition greater than 100%, 7/11 of these targets are implicated in control of pre-rRNA transcription, including MYC (Grandori et al., 2005), HEATR1/UTP10 (Gallagher et al., 2004; Prieto and McStay, 2007; Turi et al., 2018), DNTTIP2/TdIF2 (Koiwai et al., 2011), SUPT6H (Engel et al., 2015), SUPT5H (Farley-Barnes et al., 2018), EIF4A3/DDX48 (Zhang et al., 2011), and POLR2E (Goodfellow and Zomerdijk, 2013). Overall, 12/58 factors with a significant inhibition of 5-EU incorporation have been implicated in transcription, also including the RNAP1 initiation factor RRN3 (Bodem et al., 2000; Moorefield et al., 2000; Miller et al., 2001), two other t-UTPs, NOL11/UTP8 (Freed et al., 2012) and UTP4 (Freed and Baserga, 2010), and the proteins MDN1, a pre-60S assembly factor, and KIF11, a mitotic kinesin essential for RB (Ogawa et al., 2021).

Pre-rRNA processing and modification factors comprised a sizeable subset of factors with significant 5-EU mean percent inhibition. In total, 19/58 factors that inhibited nucleolar rRNA biogenesis were critical for processing, including the t-UTPs HEATR1/UTP10, NOL11/UTP8, and UTP4 (Gallagher et al., 2004; Prieto and McStay, 2007; Freed and Baserga, 2010; Freed et al., 2012; Turi et al., 2018), the C/D box snoRNP scaffolds NOP56 and NOP58 (Watkins and Bohnsack, 2012), as well as other processing factors including DNTTIP2 (Tafforeau et al., 2013), WBP11 (Carbon and Mungall, 2021), MDN1 (Raman et al., 2016), ESF1 (Tafforeau et al., 2013; Chen et al., 2018), BCCIP (Ye et al., 2020), RPP30 (Stolc and Altman, 1997; Tafforeau et al., 2013), EXOSC9 (Muller et al., 2020), NUMA1 (Farley-Barnes et al., 2018), MPHOSPH10 (Westendorf et al., 1998), TRMT112 (Zorbas et al., 2015), UTP20 (Wang et al., 2007), and NUDT16 (Ghosh et al., 2004). In addition, nucleolar rRNA biogenesis was moderately inhibited for the pre-rRNA modification factors TRMT112 (Zorbas et al., 2015), RPUSD2 (Carbon and Mungall, 2021), and NOLC1 (Yang et al., 2000; Werner et al., 2015). Notably, factors involved in transcription had a higher mean percent inhibition than factors involved in processing (83.1% inhibition v. 74.9% inhibition, n = 15 v. n = 22); factors involved in both transcription and processing had a mean percent inhibition of 99.0% (n = 6).

We also noted significant percent inhibition averages for 28 RPs from both the 40S and 60S subunits. Almost all RPs are essential for pre-rRNA biogenesis in the yeast *Saccharomyces cerevisiae* (Ferreira-Cerca et al., 2005; Poll et al., 2009) and in human cells (O’Donohue et al., 2010; Nicolas et al., 2016), compatible with a concomitant observed decrease in nucleolar 5-EU signal following their depletion.

Furthermore, of the 10 factors that had a mean percent inhibition value under 50%, five factors were either inhibitors of pre-rRNA transcription, including SUV39H1 (Murayama et al., 2008) and MAF1 (Upadhya et al., 2002; Bonhoure et al., 2020), mitochondrial ribosome biogenesis factors, including METTL15 (Van Haute et al., 2019; Chen et al., 2020) and MPV17L2 (Dalla Rosa et al., 2014), or ribosome recycling factors involved in translation, namely ABCE1 (Zhu et al., 2020).

Surprisingly, the other five RB factors with a mean percent inhibition less than 50% are well-appreciated for playing roles in pre-rRNA transcription, including POLR1D (Russell and Zomerdijk, 2006), TAF1D (Gorski et al., 2007), and TTF1 (Evers and Grummt, 1995), and in pre-rRNA processing, including NOL8 (Sekiguchi et al., 2006) and XRCC5/Ku86, which also aids TTF1 during RNAP1 termination (Wallisch et al., 2002; Shao et al., 2020). It is possible that these factors were not significantly depleted following transfection, or that, within our timeframe, the 5-EU assay cannot detect a significant change in nucleolar RNA levels as a result of non-concordant changes in both pre-rRNA transcription and stability.

Although our nucleolar 5-EU assay accurately reported the interruption of nucleolar rRNA biogenesis for the vast majority of RB factors studied, we note the following considerations and caveats regarding our method and results. First, nucleolar 5-EU incorporation can be affected by changes in one or more RB subprocesses including pre-rRNA transcription, processing, modification, and binding by RPs, which all occur co-geographically within the nucleolus. Therefore, additional mechanistic assays may be necessary to precisely define how an experimental treatment alters RB following the observation of a 5-EU defect. A treatment, like 72 h siRNA-mediated depletion of cultured human cells as we have done here, may also have opposing, compensatory effects on multiple RB subprocesses, leading to an artificially low percent inhibition and a false negative result. More broadly, as with any HTS study using RNAi-mediated target depletion, off-target effects or inefficient on-target depletion could lead to false positive or false negative results, respectively (Echeverri et al., 2006; Ou et al., 2012). Second, we have empirically defined a 5-EU significance cutoff of 50% inhibition because it minimizes the number of incorrectly classified RB factors. However, it is still unclear if there is a more stringent percent inhibition cutoff that would correspond strictly to RB factors regulating RNAP1 transcriptional activity, or cutoffs for other RB subprocesses. Future studies may elucidate the relationship between the roles of a given RB factor and the 5-EU percent inhibition value observed upon its depletion. Finally, close attention must be paid to the accuracy of nucleolar segmentation if median nucleolar 5-EU is being quantified; the maximum nuclear 5-EU metric can be used if treatment causes significant FBL dissociation, as we have observed with BMH-21.

Our miniaturized 5-EU assay enables direct quantification of nucleolar rRNA biogenesis in high-throughput, providing clearer insight into how targets modulate RB and improving upon previous HTS techniques for studying nucleolar function. The 5-EU assay is also compatible with our previously published assay for nucleolar number (Farley-Barnes et al., 2018), and is likely to be compatible with other high-content assays for ribosome biogenesis that monitor nucleolar architecture by co-staining for nucleolar proteins (He et al., 2018; Stamatopoulou et al., 2018). By extending the dimensionality and specificity of current state-of-the-art assays which indirectly track nucleolar function, our 5-EU assay will permit researchers to focus on the most promising screen candidates earlier, thereby increasing the efficiency of RB-directed screening campaigns. We anticipate that the miniaturized 5-EU assay will expedite the identification and definition of novel regulators of RB in basic or translational studies of nucleolar function.

## Materials and Methods

### Cell lines and culture conditions

Human MCF10A breast epithelial cells (ATCC CRL-10317) were cultured in DMEM/F-12 (Gibco 11330032) with 5% horse serum (Gibco 16050122), 10 μg/mL insulin (MilliporeSigma I1882), 0.5 μg/mL hydrocortisone (MilliporeSigma H0135), 20 ng/mL epidermal growth factor (Peprotech AF-100-15), and 100 ng/mL cholera toxin (MilliporeSigma C8052). Cells were incubated at 37 °C in a humidified atmosphere with 5% CO2.

### RNAi depletion by reverse-transfection

RNAi depletion was conducted in MCF10A cells as previously reported (Farley-Barnes et al., 2018; Ogawa et al., 2021). MCF10A cells were reverse-transfected into an arrayed 384-well plate library containing small interfering RNA (siRNA) constructs (Horizon Discovery, see **Supplementary Table 2**). Assay-ready plates containing 10 μL of 100 nM ON-TARGET siRNAs resuspended in 1X siRNA buffer (Horizon Discovery B-002000-UB-100) were prepared from master library 384-well plates (Horizon Discovery, 0.1 nmol scale) and stored at −80 C. Plates were thawed at room temperature for 30 min and briefly centrifuged at 300 RPM. siRNA controls (**Supplementary Table 2**) were freshly diluted in 1X siRNA buffer to 100 nM from a 50 μM frozen stock, and 10 μL of 100 nM control siRNAs were manually pipetted into the assay-ready plates. To each well, 10 μL of a 1:100 (v/v) RNAiMAX:OptiMEM solution was added (Invitrogen 13778-150, Gibco 31985070), after which the plates were briefly centrifuged at 300 RPM and incubated at room temperature for 30 min. MCF10A cells at 70%-80% confluency were trypsinized for 15 min with 0.05% trypsin (Gibco 25300054), resuspended in culture media, counted with a hemacytometer, and diluted in culture medium to a density of 100,000 cells/mL. Thirty μL of cells were dispensed into assay plates using a Multidrop Combi Reagent Dispenser (Thermo Scientific), to achieve a seeding density of 3000 cells/well, a final volume of 50 μL, and a final siRNA concentration of 20 nM. Seeded assay plates were briefly centrifuged at 300 RPM and incubated at 37 C for 72 h. Ribosome biogenesis factors were screened in triplicate.

### BMH-21 treatment and 5-ethynyl uridine incorporation

BMH-21 (MilliporeSigma SML1183) was resuspended in DMSO to a working concentration of 50 μM (50X) and stored at −20 C. 5-ethynyl uridine (5-EU, ClickChemistryTools 1261-100) was resuspended in ddH_2_O from powder to a working concentration of 50 mM (50X) and stored at −20 C. For BMH-21 treatment, reverse-transfected assay plates were treated 15 min before the end of the 72 h RNAi depletion period. One μL of 50 μM BMH-21 was manually added directly to 50 μL medium in the appropriate wells of the assay plates, which were then briefly centrifuged at 300 RPM and incubated for 15 min before 5-EU incorporation and for the remaining 1 h 5-EU treatment period. For 5-EU incorporation into nascent RNA, reverse-transfected assay plates were treated for 1 h after the end of the 72 h RNAi depletion period. One μL of 50 mM 5-EU was manually added directly to 50 μL medium in each well of the assay plates, which were then briefly centrifuged at 300 RPM and incubated for 1 h.

### Immunofluorescent staining and click fluorophore labeling

After 5-EU incorporation, cells were gently washed with 30 μL of PBS and fixed with 1% (v/v) paraformaldehyde (Electron Microscopy Sciences 15710-S) diluted in PBS at room temperature for 20 min. Cells were washed twice with 20 μL wash buffer consisting of PBS with 0.05% (v/v) TWEEN 20 (MilliporeSigma P1379), then permeabilized with 20 μL of 0.5% (v/v) Triton X-100 in PBS for 5 min. Cells were washed twice with 20 μL wash buffer and incubated with 20 μL of blocking buffer consisting of 10% (v/v) FBS (MilliporeSigma F0926) diluted in PBS for 1 h at room temperature. FBL primary antibody solution was prepared by diluting supernatant from the 72B9 hybridoma line (Reimer et al., 1987) at 1:500 or 1:250 (v/v) in blocking buffer. After blocking, cells were incubated with 20 μL FBL primary antibody solution for 2 h at room temperature. Cells were washed twice with 20 μL wash buffer and incubated with 20 μL secondary antibody solution, consisting of 1:1000 (v/v) goat anti-mouse AlexaFluor 647 (Invitrogen A-21236) and 3 μg/mL Hoechst 33342 dye in blocking buffer, for 1 h in the dark at room temperature. Immediately before the end of the secondary antibody incubation period, the click reaction cocktail was prepared in PBS by combining 5 μM AFDye 488 azide (ClickChemistryTools 1275-5), 0.5 mg/mL CuSO_4_ (Acros Organics 197730010), and 20 mg/mL freshly-resuspended sodium ascorbate (Alfa Aesar A15613). Cells were washed twice with 20 μL wash buffer, then treated with 20 μL of click reaction cocktail for 30 min in the dark at room temperature. Cells were washed twice with 20 μL wash buffer, and soaked in 20 μL PBS containing 3 μg/mL Hoechst 33342 dye for 30 min in the dark at room temperature to dissociate excess AFDye 488 azide. Cells were washed twice with 20 μL wash buffer and 40 μL of PBS was added to each well before high-content imaging.

### High-content imaging

Stained assay plates were imaged with a GE Healthcare IN Cell Analyzer 2200. Fields of view were acquired at 20X magnification with 2×2 pixel binning (665.63 μm^2^) at 16-bit depth using Cy5, DAPI, and FITC channels for FBL, Hoechst, and 5-EU staining, respectively. Laser autofocus was used to automatically determine imaging Z-height. For publication, images were cropped, merged, and labeled with scale bars using ImageJ 1.53i (Schneider et al., 2012).

### CellProfiler pipeline and data analysis

Image analysis was conducted with a custom pipeline for CellProfiler 3.1.9 (Carpenter et al., 2006; McQuin et al., 2018) (**Supplemental File 1**). Briefly, nuclei and nucleoli objects were segmented from DAPI and Cy5 channels, respectively, using global two-class Otsu thresholding. For both object classes, 5-EU intensity was measured from FITC images. Object-level normalized 5-EU intensity metrics including maximum, mean, median, and standard deviation were calculated by CellProfiler. Raw CellProfiler output CSV files including plate metadata were imported into and analyzed with JMP Pro 15.2.0 (SAS Institute). Per-well averages were computed for each 5-EU metric. For each plate, aggregate control well data were used to calculate signal-to-background (S/B) and Z’ factor screening statistics. Nucleolar RNA percent inhibition values were calculated for each well as follows:

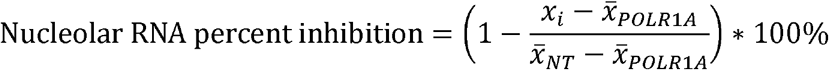

where *x* is the average 5-EU metric value over all objects in a well, *x_i_*, is the well metric value for a non-control well, 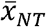 and 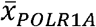 are averages of all NT or POLR1A control well metric values respectively. Plate-adjusted percent inhibition values were calculated for non-control wells by subtracting the plate’s median NT percent inhibition value from each non-control well percent inhibition (Zhang, 2011). Optimization data were graphed in JMP. Triplicate data from the ribosome biogenesis factor screen were averaged in JMP and graphed with GraphPad Prism 8 (GraphPad Software).

## Acknowledgements and Funding Sources

We thank the members of the laboratory of S.J.B. and S. Carter for insightful information, questions, and comments throughout the manuscript writing process. We acknowledge the use of CellProfiler for image analysis (http://www.cellprofiler.org/). This work was supported by the following grants from the National Institutes of Health (NIH): 1R35GM131687 (to S.J.B.), 1F31DE030332 (to M.A.M.), and T32GM007223 (to C.J.B., M.A.M., and S.J.B.).

---

**Supplementary Figure 1.**
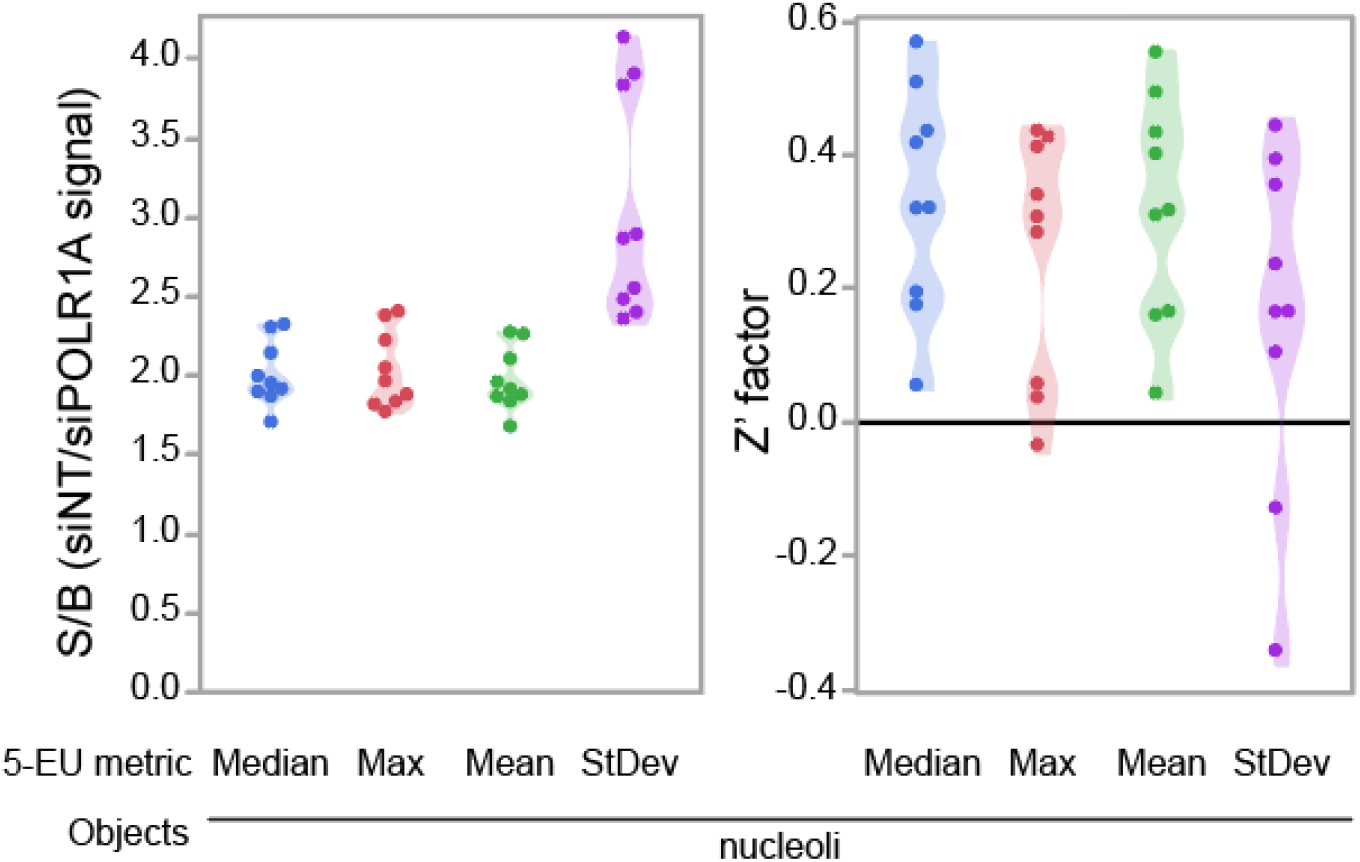
Control S/B and Z’ factor values for validation assay replicates. Control S/B is calculated as the ratio of mean siNT-treated 5-EU signal divided by mean siPOLR1A-treated 5-EU signal. Median (blue), maximum (red), mean (green), or standard deviation (purple) of nucleolar 5-EU signal. n = 9 replicates.

