## Supplemental File 1 for "A high-throughput assay for directly monitoring nucleolar rRNA biogenesis"

CellProfiler Pipeline: http://www.cellprofiler.org
Version:4
DateRevision:319
GitHash:
ModuleCount:8
HasImagePlaneDetails:False

Images:[module_num:1|svn_version:\'Unknown\'|variable_revision_number:2|show_window:False|notes:\x5B\'To begin creating your project, use the Images module to compile a list of files and/or folders that you want to analyze. You can also specify a set of rules to include only the desired files in your selected folders.\'\x5D|batch_state:array(\x5B\x5D, dtype=uint8)|enabled:True|wants_pause:False]
    :\xff\xfe
    Filter images?:\xff\xfeC\x00u\x00s\x00t\x00o\x00m\x00
    Select the rule criteria:\xff\xfea\x00n\x00d\x00 \x00(\x00e\x00x\x00t\x00e\x00n\x00s\x00i\x00o\x00n\x00 \x00d\x00o\x00e\x00s\x00 \x00i\x00s\x00t\x00i\x00f\x00)\x00

Metadata:[module_num:2|svn_version:\'Unknown\'|variable_revision_number:5|show_window:False|notes:\x5B\'The Metadata module optionally allows you to extract information describing your images (i.e, metadata) which will be stored along with your measurements. This information can be contained in the file name and/or location, or in an external file.\'\x5D|batch_state:array(\x5B\x5D, dtype=uint8)|enabled:True|wants_pause:False]
    Extract metadata?:\xff\xfeY\x00e\x00s\x00
    Metadata data type:\xff\xfeT\x00e\x00x\x00t\x00
    Metadata types:\xff\xfe{\x00}\x00
    Extraction method count:\xff\xfe1\x00
    Metadata extraction method:\xff\xfeE\x00x\x00t\x00r\x00a\x00c\x00t\x00 \x00f\x00r\x00o\x00m\x00 \x00f\x00i\x00l\x00e\x00/\x00f\x00o\x00l\x00d\x00e\x00r\x00 \x00n\x00a\x00m\x00e\x00s\x00
    Metadata source:\xff\xfeF\x00i\x00l\x00e\x00 \x00n\x00a\x00m\x00e\x00
    Regular expression to extract from file name:\xff\xfe
    Regular expression to extract from folder name:\xff\xfe(\x00?\x00P\x00<\x00D\x00a\x00t\x00e\x00>\x00\x5B\x000\x00-\x009\x00\x5D\x00{\x004\x00}\x00_\x00\x5B\x000\x00-\x009\x00\x5D\x00{\x002\x00}\x00_\x00\x5B\x000\x00-\x009\x00\x5D\x00{\x002\x00}\x00)\x00$\x00
    Extract metadata from:\xff\xfeA\x00l\x00l\x00 \x00i\x00m\x00a\x00g\x00e\x00s\x00
    Select the filtering criteria:\xff\xfea\x00n\x00d\x00 \x00(\x00f\x00i\x00l\x00e\x00 \x00d\x00o\x00e\x00s\x00 \x00c\x00o\x00n\x00t\x00a\x00i\x00n\x00 \x00"\x00"\x00)\x00
    Metadata file location:\xff\xfeE\x00l\x00s\x00e\x00w\x00h\x00e\x00r\x00e\x00.\x00.\x00.\x00\x7C\x00
    Match file and image metadata:\xff\xfe\x5B\x00\x5D\x00
    Use case insensitive matching?:\xff\xfeN\x00o\x00
    Metadata file name:\xff\xfe

NamesAndTypes:[module_num:3|svn_version:\'Unknown\'|variable_revision_number:8|show_window:False|notes:\x5B\'The NamesAndTypes module allows you to assign a meaningful name to each image by which other modules will refer to it.\'\x5D|batch_state:array(\x5B\x5D, dtype=uint8)|enabled:True|wants_pause:False]
    Assign a name to:\xff\xfeI\x00m\x00a\x00g\x00e\x00s\x00 \x00m\x00a\x00t\x00c\x00h\x00i\x00n\x00g\x00 \x00r\x00u\x00l\x00e\x00s\x00
    Select the image type:\xff\xfeG\x00r\x00a\x00y\x00s\x00c\x00a\x00l\x00e\x00 \x00i\x00m\x00a\x00g\x00e\x00
    Name to assign these images:\xff\xfeD\x00A\x00P\x00I\x00
    Match metadata:\xff\xfe\x5B\x00\x5D\x00
    Image set matching method:\xff\xfeO\x00r\x00d\x00e\x00r\x00
    Set intensity range from:\xff\xfeI\x00m\x00a\x00g\x00e\x00 \x00b\x00i\x00t\x00-\x00d\x00e\x00p\x00t\x00h\x00
    Assignments count:\xff\xfe3\x00
    Single images count:\xff\xfe0\x00
    Maximum intensity:\xff\xfe2\x005\x005\x00.\x000\x00
    Process as 3D?:\xff\xfeN\x00o\x00
    Relative pixel spacing in X:\xff\xfe1\x00.\x000\x00
    Relative pixel spacing in Y:\xff\xfe1\x00.\x000\x00
    Relative pixel spacing in Z:\xff\xfe1\x00.\x000\x00
    Select the rule criteria:\xff\xfea\x00n\x00d\x00 \x00(\x00f\x00i\x00l\x00e\x00 \x00d\x00o\x00e\x00s\x00 \x00c\x00o\x00n\x00t\x00a\x00i\x00n\x00 \x00"\x00D\x00A\x00P\x00I\x00"\x00)\x00
    Name to assign these images:\xff\xfeD\x00N\x00A\x00
    Name to assign these objects:\xff\xfeC\x00e\x00l\x00l\x00
    Select the image type:\xff\xfeG\x00r\x00a\x00y\x00s\x00c\x00a\x00l\x00e\x00 \x00i\x00m\x00a\x00g\x00e\x00
    Set intensity range from:\xff\xfeI\x00m\x00a\x00g\x00e\x00 \x00b\x00i\x00t\x00-\x00d\x00e\x00p\x00t\x00h\x00
    Maximum intensity:\xff\xfe2\x005\x005\x00.\x000\x00
    Select the rule criteria:\xff\xfea\x00n\x00d\x00 \x00(\x00f\x00i\x00l\x00e\x00 \x00d\x00o\x00e\x00s\x00 \x00c\x00o\x00n\x00t\x00a\x00i\x00n\x00 \x00"\x00C\x00y\x005\x00"\x00)\x00
    Name to assign these images:\xff\xfeF\x00i\x00b\x00r\x00i\x00l\x00l\x00a\x00r\x00i\x00n\x00
    Name to assign these objects:\xff\xfeN\x00u\x00c\x00l\x00e\x00u\x00s\x00
    Select the image type:\xff\xfeG\x00r\x00a\x00y\x00s\x00c\x00a\x00l\x00e\x00 \x00i\x00m\x00a\x00g\x00e\x00
    Set intensity range from:\xff\xfeI\x00m\x00a\x00g\x00e\x00 \x00b\x00i\x00t\x00-\x00d\x00e\x00p\x00t\x00h\x00
    Maximum intensity:\xff\xfe2\x005\x005\x00.\x000\x00
    Select the rule criteria:\xff\xfea\x00n\x00d\x00 \x00(\x00f\x00i\x00l\x00e\x00 \x00d\x00o\x00e\x00s\x00 \x00c\x00o\x00n\x00t\x00a\x00i\x00n\x00 \x00"\x00F\x00I\x00T\x00C\x00"\x00)\x00
    Name to assign these images:\xff\xfeO\x00P\x00P\x00_\x005\x00E\x00U\x00
    Name to assign these objects:\xff\xfeC\x00y\x00t\x00o\x00p\x00l\x00a\x00s\x00m\x00
    Select the image type:\xff\xfeG\x00r\x00a\x00y\x00s\x00c\x00a\x00l\x00e\x00 \x00i\x00m\x00a\x00g\x00e\x00
    Set intensity range from:\xff\xfeI\x00m\x00a\x00g\x00e\x00 \x00m\x00e\x00t\x00a\x00d\x00a\x00t\x00a\x00
    Maximum intensity:\xff\xfe2\x005\x005\x00.\x000\x00

Groups:[module_num:4|svn_version:\'Unknown\'|variable_revision_number:2|show_window:False|notes:\x5B\'The Groups module optionally allows you to split your list of images into image subsets (groups) which will be processed independently of each other. Examples of groupings include screening batches, microtiter plates, time-lapse movies, etc.\'\x5D|batch_state:array(\x5B\x5D, dtype=uint8)|enabled:True|wants_pause:False]
    Do you want to group your images?:\xff\xfeN\x00o\x00
    grouping metadata count:\xff\xfe1\x00
    Metadata category:\xff\xfeN\x00o\x00n\x00e\x00

IdentifyPrimaryObjects:[module_num:5|svn_version:\'Unknown\'|variable_revision_number:13|show_window:False|notes:\x5B\'Identify Nuclei from Hoechst/DAPI signal\'\x5D|batch_state:array(\x5B\x5D, dtype=uint8)|enabled:True|wants_pause:False]
    Select the input image:\xff\xfeD\x00N\x00A\x00
    Name the primary objects to be identified:\xff\xfeN\x00u\x00c\x00l\x00e\x00i\x00
    Typical diameter of objects, in pixel units (Min,Max):\xff\xfe1\x000\x00,\x005\x000\x00
    Discard objects outside the diameter range?:\xff\xfeY\x00e\x00s\x00
    Discard objects touching the border of the image?:\xff\xfeY\x00e\x00s\x00
    Method to distinguish clumped objects:\xff\xfeS\x00h\x00a\x00p\x00e\x00
    Method to draw dividing lines between clumped objects:\xff\xfeS\x00h\x00a\x00p\x00e\x00
    Size of smoothing filter:\xff\xfe1\x000\x00
    Suppress local maxima that are closer than this minimum allowed distance:\xff\xfe7\x00.\x000\x00
    Speed up by using lower-resolution image to find local maxima?:\xff\xfeY\x00e\x00s\x00
    Fill holes in identified objects?:\xff\xfeA\x00f\x00t\x00e\x00r\x00 \x00b\x00o\x00t\x00h\x00 \x00t\x00h\x00r\x00e\x00s\x00h\x00o\x00l\x00d\x00i\x00n\x00g\x00 \x00a\x00n\x00d\x00 \x00d\x00e\x00c\x00l\x00u\x00m\x00p\x00i\x00n\x00g\x00
    Automatically calculate size of smoothing filter for declumping?:\xff\xfeY\x00e\x00s\x00
    Automatically calculate minimum allowed distance between local maxima?:\xff\xfeY\x00e\x00s\x00
    Handling of objects if excessive number of objects identified:\xff\xfeC\x00o\x00n\x00t\x00i\x00n\x00u\x00e\x00
    Maximum number of objects:\xff\xfe5\x000\x000\x00
    Use advanced settings?:\xff\xfeY\x00e\x00s\x00
    Threshold setting version:\xff\xfe1\x000\x00
    Threshold strategy:\xff\xfeG\x00l\x00o\x00b\x00a\x00l\x00
    Thresholding method:\xff\xfeO\x00t\x00s\x00u\x00
    Threshold smoothing scale:\xff\xfe1\x00
    Threshold correction factor:\xff\xfe1\x00.\x000\x00
    Lower and upper bounds on threshold:\xff\xfe0\x00.\x000\x00,\x001\x00.\x000\x00
    Manual threshold:\xff\xfe0\x00.\x000\x00
    Select the measurement to threshold with:\xff\xfeN\x00o\x00n\x00e\x00
    Two-class or three-class thresholding?:\xff\xfeT\x00w\x00o\x00 \x00c\x00l\x00a\x00s\x00s\x00e\x00s\x00
    Assign pixels in the middle intensity class to the foreground or the background?:\xff\xfeF\x00o\x00r\x00e\x00g\x00r\x00o\x00u\x00n\x00d\x00
    Size of adaptive window:\xff\xfe5\x000\x00
    Lower outlier fraction:\xff\xfe0\x00.\x000\x005\x00
    Upper outlier fraction:\xff\xfe0\x00.\x000\x005\x00
    Averaging method:\xff\xfeM\x00e\x00a\x00n\x00
    Variance method:\xff\xfeS\x00t\x00a\x00n\x00d\x00a\x00r\x00d\x00 \x00d\x00e\x00v\x00i\x00a\x00t\x00i\x00o\x00n\x00
    # of deviations:\xff\xfe2\x00.\x000\x00
    Thresholding method:\xff\xfeO\x00t\x00s\x00u\x00

IdentifyPrimaryObjects:[module_num:6|svn_version:\'Unknown\'|variable_revision_number:13|show_window:False|notes:\x5B\'Identify Nucleoli from FBL/Cy5 signal\'\x5D|batch_state:array(\x5B\x5D, dtype=uint8)|enabled:True|wants_pause:False]
    Select the input image:\xff\xfeF\x00i\x00b\x00r\x00i\x00l\x00l\x00a\x00r\x00i\x00n\x00
    Name the primary objects to be identified:\xff\xfeN\x00u\x00c\x00l\x00e\x00o\x00l\x00i\x00
    Typical diameter of objects, in pixel units (Min,Max):\xff\xfe2\x00,\x002\x000\x00
    Discard objects outside the diameter range?:\xff\xfeY\x00e\x00s\x00
    Discard objects touching the border of the image?:\xff\xfeY\x00e\x00s\x00
    Method to distinguish clumped objects:\xff\xfeS\x00h\x00a\x00p\x00e\x00
    Method to draw dividing lines between clumped objects:\xff\xfeS\x00h\x00a\x00p\x00e\x00
    Size of smoothing filter:\xff\xfe1\x00
    Suppress local maxima that are closer than this minimum allowed distance:\xff\xfe1\x00.\x000\x00
    Speed up by using lower-resolution image to find local maxima?:\xff\xfeY\x00e\x00s\x00
    Fill holes in identified objects?:\xff\xfeA\x00f\x00t\x00e\x00r\x00 \x00b\x00o\x00t\x00h\x00 \x00t\x00h\x00r\x00e\x00s\x00h\x00o\x00l\x00d\x00i\x00n\x00g\x00 \x00a\x00n\x00d\x00 \x00d\x00e\x00c\x00l\x00u\x00m\x00p\x00i\x00n\x00g\x00
    Automatically calculate size of smoothing filter for declumping?:\xff\xfeY\x00e\x00s\x00
    Automatically calculate minimum allowed distance between local maxima?:\xff\xfeY\x00e\x00s\x00
    Handling of objects if excessive number of objects identified:\xff\xfeC\x00o\x00n\x00t\x00i\x00n\x00u\x00e\x00
    Maximum number of objects:\xff\xfe5\x000\x000\x00
    Use advanced settings?:\xff\xfeY\x00e\x00s\x00
    Threshold setting version:\xff\xfe1\x000\x00
    Threshold strategy:\xff\xfeG\x00l\x00o\x00b\x00a\x00l\x00
    Thresholding method:\xff\xfeO\x00t\x00s\x00u\x00
    Threshold smoothing scale:\xff\xfe1\x00
    Threshold correction factor:\xff\xfe1\x00.\x002\x00
    Lower and upper bounds on threshold:\xff\xfe0\x00.\x000\x00,\x001\x00.\x000\x00
    Manual threshold:\xff\xfe0\x00.\x000\x00
    Select the measurement to threshold with:\xff\xfeN\x00o\x00n\x00e\x00
    Two-class or three-class thresholding?:\xff\xfeT\x00w\x00o\x00 \x00c\x00l\x00a\x00s\x00s\x00e\x00s\x00
    Assign pixels in the middle intensity class to the foreground or the background?:\xff\xfeB\x00a\x00c\x00k\x00g\x00r\x00o\x00u\x00n\x00d\x00
    Size of adaptive window:\xff\xfe5\x000\x00
    Lower outlier fraction:\xff\xfe0\x00.\x000\x005\x00
    Upper outlier fraction:\xff\xfe0\x00.\x000\x005\x00
    Averaging method:\xff\xfeM\x00e\x00a\x00n\x00
    Variance method:\xff\xfeS\x00t\x00a\x00n\x00d\x00a\x00r\x00d\x00 \x00d\x00e\x00v\x00i\x00a\x00t\x00i\x00o\x00n\x00
    # of deviations:\xff\xfe2\x00.\x000\x00
    Thresholding method:\xff\xfeO\x00t\x00s\x00u\x00

MeasureObjectIntensity:[module_num:7|svn_version:\'Unknown\'|variable_revision_number:3|show_window:False|notes:\x5B\'Measure DNA from DAPI and 5-EU from FITC signal -- measure in both whole nuclei and within nucleoli; select measurements in ExportToSpreadhseet\'\x5D|batch_state:array(\x5B\x5D, dtype=uint8)|enabled:True|wants_pause:False]
    Hidden:\xff\xfe1\x00
    Select an image to measure:\xff\xfeO\x00P\x00P\x00_\x005\x00E\x00U\x00
    Select objects to measure:\xff\xfeN\x00u\x00c\x00l\x00e\x00i\x00
    Select objects to measure:\xff\xfeN\x00u\x00c\x00l\x00e\x00o\x00l\x00i\x00

ExportToSpreadsheet:[module_num:8|svn_version:\'Unknown\'|variable_revision_number:12|show_window:False|notes:\x5B\'Export output to spreadsheet -- need to change output file location, sub-folder, and filename prefix for each run\', \'Can adjust which measurements are exported using "Press button to select measurements" (try to minimize to save file space)\', \'\', \'Read data in JMP, Excel, R, Python, etc.\'\x5D|batch_state:array(\x5B\x5D, dtype=uint8)|enabled:True|wants_pause:False]
    Select the column delimiter:\xff\xfeC\x00o\x00m\x00m\x00a\x00 \x00(\x00"\x00,\x00"\x00)\x00
    Add image metadata columns to your object data file?:\xff\xfeY\x00e\x00s\x00
    Select the measurements to export:\xff\xfeY\x00e\x00s\x00
    Calculate the per-image mean values for object measurements?:\xff\xfeN\x00o\x00
    Calculate the per-image median values for object measurements?:\xff\xfeN\x00o\x00
    Calculate the per-image standard deviation values for object measurements?:\xff\xfeN\x00o\x00
    Output file location:\xff\xfeD\x00e\x00f\x00a\x00u\x00l\x00t\x00 \x00I\x00n\x00p\x00u\x00t\x00 \x00F\x00o\x00l\x00d\x00e\x00r\x00 \x00s\x00u\x00b\x00-\x00f\x00o\x00l\x00d\x00e\x00r\x00\x7C\x00
    Create a GenePattern GCT file?:\xff\xfeN\x00o\x00
    Select source of sample row name:\xff\xfeM\x00e\x00t\x00a\x00d\x00a\x00t\x00a\x00
    Select the image to use as the identifier:\xff\xfeN\x00o\x00n\x00e\x00
    Select the metadata to use as the identifier:\xff\xfeN\x00o\x00n\x00e\x00
    Export all measurement types?:\xff\xfeY\x00e\x00s\x00
    Press button to select measurements:\xff\xfeE\x00x\x00p\x00e\x00r\x00i\x00m\x00e\x00n\x00t\x00\x7C\x00M\x00o\x00d\x00i\x00f\x00i\x00c\x00a\x00t\x00i\x00o\x00n\x00_\x00T\x00i\x00m\x00e\x00s\x00t\x00a\x00m\x00p\x00,\x00E\x00x\x00p\x00e\x00r\x00i\x00m\x00e\x00n\x00t\x00\x7C\x00P\x00i\x00p\x00e\x00l\x00i\x00n\x00e\x00_\x00P\x00i\x00p\x00e\x00l\x00i\x00n\x00e\x00,\x00E\x00x\x00p\x00e\x00r\x00i\x00m\x00e\x00n\x00t\x00\x7C\x00C\x00e\x00l\x00l\x00P\x00r\x00o\x00f\x00i\x00l\x00e\x00r\x00_\x00V\x00e\x00r\x00s\x00i\x00o\x00n\x00,\x00E\x00x\x00p\x00e\x00r\x00i\x00m\x00e\x00n\x00t\x00\x7C\x00R\x00u\x00n\x00_\x00T\x00i\x00m\x00e\x00s\x00t\x00a\x00m\x00p\x00,\x00I\x00m\x00a\x00g\x00e\x00\x7C\x00C\x00o\x00u\x00n\x00t\x00_\x00N\x00u\x00c\x00l\x00e\x00i\x00,\x00N\x00u\x00c\x00l\x00e\x00i\x00\x7C\x00I\x00n\x00t\x00e\x00n\x00s\x00i\x00t\x00y\x00_\x00M\x00a\x00x\x00I\x00n\x00t\x00e\x00n\x00s\x00i\x00t\x00y\x00_\x00O\x00P\x00P\x00_\x005\x00E\x00U\x00,\x00N\x00u\x00c\x00l\x00e\x00i\x00\x7C\x00N\x00u\x00m\x00b\x00e\x00r\x00_\x00O\x00b\x00j\x00e\x00c\x00t\x00_\x00N\x00u\x00m\x00b\x00e\x00r\x00,\x00N\x00u\x00c\x00l\x00e\x00o\x00l\x00i\x00\x7C\x00I\x00n\x00t\x00e\x00n\x00s\x00i\x00t\x00y\x00_\x00M\x00e\x00d\x00i\x00a\x00n\x00I\x00n\x00t\x00e\x00n\x00s\x00i\x00t\x00y\x00_\x00O\x00P\x00P\x00_\x005\x00E\x00U\x00
    Representation of Nan/Inf:\xff\xfeN\x00a\x00N\x00
    Add a prefix to file names?:\xff\xfeY\x00e\x00s\x00
    Filename prefix:\xff\xfe
    Overwrite existing files without warning?:\xff\xfeN\x00o\x00
    Data to export:\xff\xfeD\x00o\x00 \x00n\x00o\x00t\x00 \x00u\x00s\x00e\x00
    Combine these object measurements with those of the previous object?:\xff\xfeN\x00o\x00
    File name:\xff\xfeD\x00A\x00T\x00A\x00.\x00c\x00s\x00v\x00
    Use the object name for the file name?:\xff\xfeY\x00e\x00s\x00
